# Adaptation by copy number variation increases insecticide resistance in fall armyworms

**DOI:** 10.1101/812958

**Authors:** Kiwoong Nam, Sylvie Gimenez, Frederique Hilliou, Carlos A. Blanco, Sabine Hänniger, Anthony Bretaudeau, Fabrice Legeai, Nicolas Nègre, Emmanuelle d’Alençon

## Abstract

Insecticide resistance is a major main challenge in pest control, and understanding its genetic basis is a key topic in agricultural ecology. Detoxification genes are well-known genetic elements that play a key role in adaptation to xenobiotics. The adaptive evolution of detoxification genes by copy number variations has been interpreted as a cause of insecticide resistance. However, the same pattern can be generated by the adaptation to host-plant defense toxins as well. In this study, we tested in fall armyworms (Lepidoptera *Spodoptera frugiperda*) if adaptation by copy number variation is the cause of the increased level of insecticide resistance from two geographic populations with different levels of resistance and two strains with different host plants. Following the generation of an assembly with chromosome-sized scaffolds (N50 = 13.2Mb), we observed that these two populations show a significant allelic differentiation of copy number variations, which is not observed between strains. In particular, a locus with almost complete allelic differentiation (Fst > 0.8) includes a cluster of P450 genes, which are well-known key players in detoxification. Detoxification genes are overrepresented in the genes with copy number variations, and the observed copy number variation appears to have beneficial effects in general. From this result, we concluded that copy number variation of detoxification genes in fall armyworm*s* plays a key role in the insecticide resistance but not in the adaptation to host-plants, suggesting that the evolution of insecticide resistance may occur independently from host-plant adaptation.

## Inroduction

The emergence of insecticide resistance is one of the biggest challenges in pest control. In the USA, the resistance causes an extra cost of billions of US dollars every year^1^. The identification of genes responsible for insecticide resistance has been one of the main topics in agricultural ecology because this knowledge can be used to design effective ways of controlling pests^2^.Well-known detoxification genes include cytochrome P450s (CYPs)^3^, esterases^4^,Glutathione S-transferases (GSTs)^3^, UDP glucuronosyltransferases (UGTs)^5^, and oxidative stress genes^6^.The copy number variation (CNV) of these detoxication genes and associated positive selection have been reported from several insect pest species, including mosquitoes (*Anopheles gambiae*)^7,8^,Tobacco cutworm (*Spodoptera litura*)^9^, and fall armyworms (*S. frugiperda*)^10^.These genetic mechanisms are assumed to be associated with insecticide resistance^11^.

However, positive selection on detoxification gene itself is not necessarily a cause of insecticide resistance. Insects produce detoxification proteins to overcome plant defense toxins. Thus, the adaptive CNV of detoxification genes can be a consequence of evolutionary arms races between insects and plants^12^. As the evolutionary history of host-plant adaptation is much longer than that involving human insecticide application^1^, the vast majority of evolutionary genetic footprints of detoxification genes might be generated independently from insecticides. Therefore, the association between the CNV of detoxification genes and insecticide resistance remains elusive.

The fall armyworm (*Spodoptera frugiperda*) is one of the most damaging pest insects of different classes of crop plants due to its extreme polyphagy and strong migratory behavior. The fall armyworm is native to almost all the entire North and South American continents, and in 2016 this species invaded sub-Saharan Africa.13. Then, fall armyworms globally spread by the invasions of India^14^, South East Asia, East Asia, and Egypt (https://www.cabi.org/isc/fallarmyworm). Interestingly, this species consists of two sympatric strains, the corn strain (sfC) and the rice strains (sfR)^15–17^. The strains named after their preferred host-plants, sfCs prefers tall grasses like corn or sorghum, and the sfR prefers smaller grasses like rice or pasture grasses. The differentiated host-plant ranges imply different adaptation processes to host-plants, probably involving differentiated detoxification processes^18^. In Puerto Rico, fall armyworms show a dramatically increased level of field-evolved insecticide resistance from a wide range of insecticides compared with populations in the mainland of the USA (e.g., Monsanto strain)^19^ and Mexico^20^. The reason for this increase is not well known, but one of the possibilities is strong selective pressure by massive sprays of insecticides for corn seed production^21^.

Therefore, the fall armyworm population from Puerto Rico is an ideal model species to distinguish the adaptive role of CNVs between host-plant adaptation and insecticide resistance. If positive selection specific to Puerto Rico’s population is observed from the CNVs of detoxification genes while these CNVs are not specific to strains, then the adaptive role of CNVs in insecticide resistance is supported. Alternatively, if all identified positive selection by CNVs is specific to the host-plant strains, then the CNVs contributed predominantly to the adaptation to host-plants. To test these possibilities, we analyze the resequencing data from fall armyworm in Puerto Rico and Mississippi, including sfC and sfR for each population, together with a new high-quality chromosome-sized reference genome assembly.

## Results

### Reference genome and resequencing data

We improved the reference genome assemblies from sfC and sfR used in our previous study^22^ because the existing reference genome assembly from Pac-Bio reads is moderately fragmented (1,000 scaffolds, N50 = 900kb for sfC). We performed super-scaffolding from the genome assembly from sfC using Hi-C^23^ data and HiRise software^24^.The assembly size is 384.46Mb, which is close to the expectation by flow cytometry (396±3 Mb)^10^. The number of sequences in the improved reference genome assembly is 125, and N50 is 13.15Mb. The number of the longest super-scaffolds explaining 90% of genome assembly (L90) is 27. The scaffold lengths show a bimodal distribution, and a mode corresponding to longer sequences (>5Mb) contains 31 scaffolds (Supplementary figure 1). As the fall armyworm has 31 chromosomes in the genome, this pattern demonstrates a chromosome-sized assembly. Recently, Zhang et al. also generated a high-quality chromosome-sized assembly generated from an invasive African population (submitted to *Nature communications*). We believe that these two chromosome-sized assemblies are complementary between each other because the assemblies generated in this study and by Zhang et al. represent native and invasive populations, respectively.

Recently, Liu et al. published another new reference genome assemblies from invasive fall armyworms^25^. The assemblies are substantially larger than the expected genome size (530.77Mb - 542.42Mb), raising the possibility of misassemblies. As Liu et al. generated the genome assemblies from a natural population, a potentially high level of heterozygosity might inflate the assembly size. To evaluate the correctness of the assemblies, we performed BUSCO (Benchmarking Universal Single-Copy Ortholog) analysis^26^. Our new assembly has a higher proportion of complete BUSCO genes than the assembly by Liu et al (Table 1). In addition, our new assembly has a lower proportion of duplicated BUSCO genes (28/1658 = 1.69%) than the assemblies from Liu et al^25^ (134/1658 = 8.08% for male assembly, 97/1658 = 5.86% for female assembly). From this BUSCO analysis, we concluded that our assembly has higher accuracy than the assemblies from Liu et al^25^. In addition, our assembly has better contiguity because of a much smaller number of contigs and a much smaller L90 value than the assembly by Liu et al^25^ (Table 1).

**Table 1.** Summary statistics of reference genome assembly and the result of BUSCO analysis from the assembles used in this study and in Liu et al^25^.

| Assembly statistics | ver6 | Liu et al, Male | Liu et al, Female |
| --- | --- | --- | --- |
| Assembly size (bp) | 384,455,365 | 543,659,128 | 531,931,622 |
| Number of sequences | 125 | 21,840 | 27,258 |
| N50 (bp) | 13,151,234 | 14,162,803 | 13,967,093 |
| L50 | 13 | 16 | 17 |
| N90 (bp) | 8,473,354 | 6,440 | 5,122 |
| L90 | 27 | 3,030 | 5,612 |
| Length of gaps (bp) | 346,864 | 37,953,553 | 35,713,062 |
| Complete and single-copy BUSCOs | 1,573 | 1,442 | 1,480 |
| Complete and duplicated BUSCOs | 28 | 134 | 97 |
| Fragmented BUSCOs | 20 | 45 | 48 |
| Missing BUSCOs | 37 | 37 | 33 |

The resequencing data were obtained from a population in Mississippi (MS) and from a Puerto Rico population (PR) using Illumina sequencing technology. Then, we used mitochondrial genomes to identify strains for each individual (Supplementary figure 2). Mississippi resequencing data contains nine sfC and eight sfR, respectively. PR contains 11 sfC and four sfR, respectively. Then, we performed mapping of the resequencing data against the reference genome assembly. SNV (single nucleotide variants) and CNV were identified using GATK^27^ and CNVcaller^28^, respectively (Supplementary figure 3). For the CNVs, we used only those showing minor allele frequency of variant greater than 0.1 in order to reduce false positives. The numbers of identified CNVs are 41,645, and the average size of the CNV unit is 1501.14bp (800bp - 151kb). For SNVs, we identified 16,341,783 after strict filtering.

### CNV according to Geographical differentiation

We performed principal component analysis (PCA) to infer the genetic relationship among individuals from each of CNVs and SNVs. The result from CNVs shows a clear grouping according to the geographical population, whereas the grouping according to the strain is not observed (Fig 1). SNVs also showed a weak pattern of grouping according to the geographical populations, but not according to strains. The PCA result shows that PR has a much smaller variation of CNV than the MS, while SNVs do not show such a pattern. This result is in line with prevalent natural selection by CNVs specific to PR, which is not observed from SNVs.

**Figure 1.**
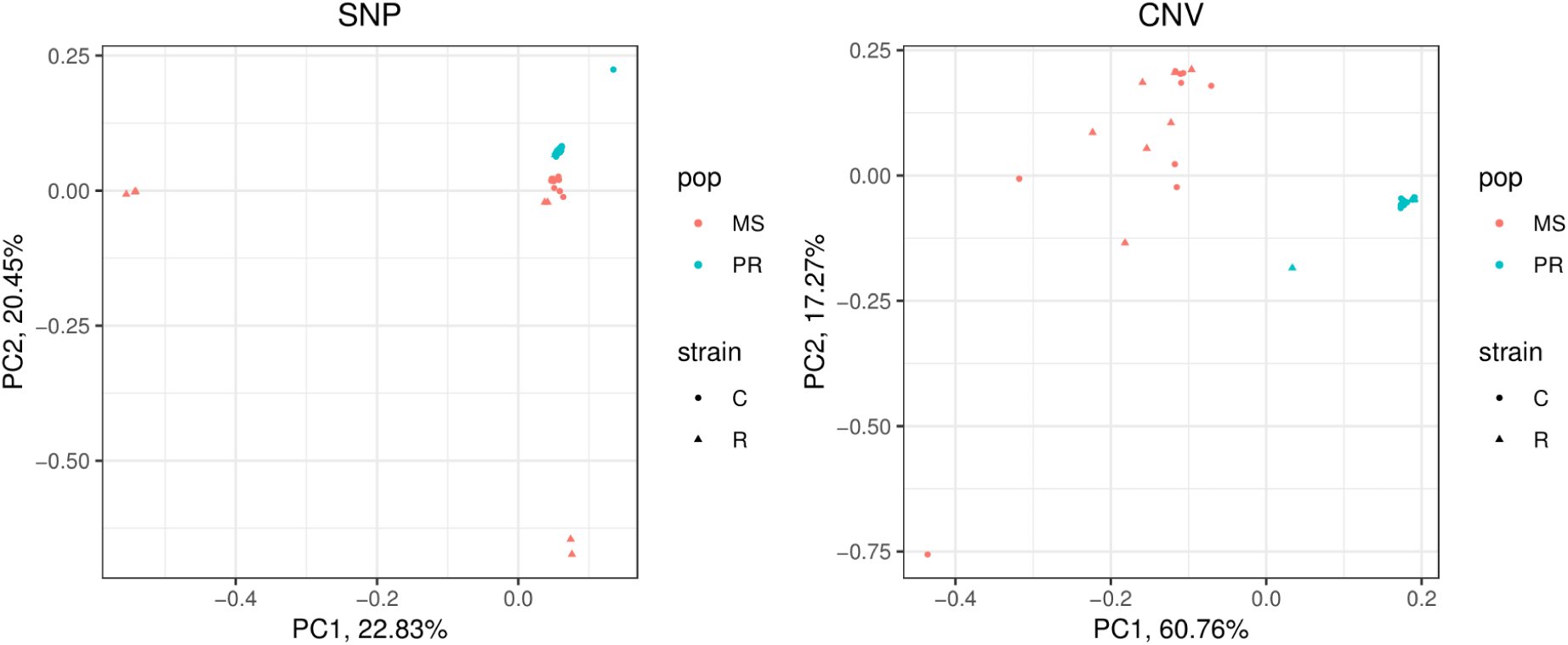
Principal component analysis from SNP (left) and CNV (right). MS and PR represent samples from Mississippi and Puerto Rico, respectively. And the C and R represent sfC and sfR, respectively.

Fst calculated from CNVs and SNVs between PR and MS is 0.142 and 0.0163, respectively, and both Fst values are significantly greater than the expectation based on random grouping (p < 0.001 for both CNV and SNV). This result implies that both CNVs and SNVs show significant genetic differentiation between PR and MS, with a much greater extent for CNV, in line with the PCA results (Fig 1). Fst calculated from CNV between strains is not significantly greater than the expectation based on random grouping (Fst = 0.0019, p = 0.262), while Fst calculated from SNV is very low but significantly higher than the expectation by random grouping (Fst = 0.0093, p < 0.001). This result shows that the genomic distribution of CNVs has been (re)shaped by evolutionary forces that are specific to geographic population(s), but not specific to host-plants.

However, Fst between sfC and sfR may not represent the level of genetic differentiation between the strains if hybrids exist in the samples we used in this study. Therefore, we test if potential hybrids affect the calculated Fst calculation between strains significantly. All the samples we used in this study are females, which has ZW sex chromosomes (please note that males have ZZ chromosomes). The Z chromosomes in females were inherited from the fathers, whereas the mitochondrial genomes were inherited from the mothers. Therefore, female Z chromosome and mitochondrial genomes are inherited from different strains in hybrids. In this study, the strain was identified from mitochondrial genomes. If a significant proportion of these samples are hybrids, Z chromosomes will have lower Fst between strains than nuclear genomes (see Supplementary figure 4 for more detail). We performed blasting of Z-linked TPI genes^29^ to identify Z chromosome in the assembly, and we observed only a single blast-hit, at Scaffold 66. This scaffold is 21,694,391bp in length. This Z chromosome has higher Fst than autosomes (Fst for Z chromosome and autosome is 0.024 and 0.0088, respectively, p = 0.0085; bootstrapping test). Therefore, it is unlikely that hybrids affect the results significantly.

Then, we inferred geographic population-specific positive selection from the loci with nearly complete allelic differentiation of CNVs between PR and MS (Fst > 0.8). In total, we identified seven positively selected loci (Fig 2A). Among these loci, the unit of the CNV at scaffold 17 is a gene cluster, which is composed of 12 CYP9A genes and two alcohol dehydrogenase (ADH) genes (Fig 2B). This gene cluster was also observed in our previous study based on BAC from sfC^30^, which has been raised in our insectarium. In PR all 30 alleles have two copies of this unit, while 28 and six alleles in the MS have one and two copies, respectively. The presence of a single haplotype of CNVs in PR and of two haplotypes of CNVs in the MS imply the existence of positive selection that is specific to PR because positive selection fixes only a single haplotype in a population. The overexpression of CYP9A genes upon the treatment of insecticides was observed from the fall armyworm^31^, Beet armyworm (*S.exigua)*^32^, and Tobacco cutworm (*S.litura*)^33^, and smaller tea tortrix (Adoxophyes honmai)^34^, supporting the association between CYP9A and insecticide resistance. The presence of ADH gene cluster was also found in a previous study^30^. In Lepidopteran *Helicoverpa armigera*, ADH5 binds a promoter of CYP6B6 in the response of xenobiotic 2-tridecanone. Deciphering the functions of CYP and ADH genes from scaffold 17 will contribute to the understanding of insecticide resistance in PR.

**Figure 2.**
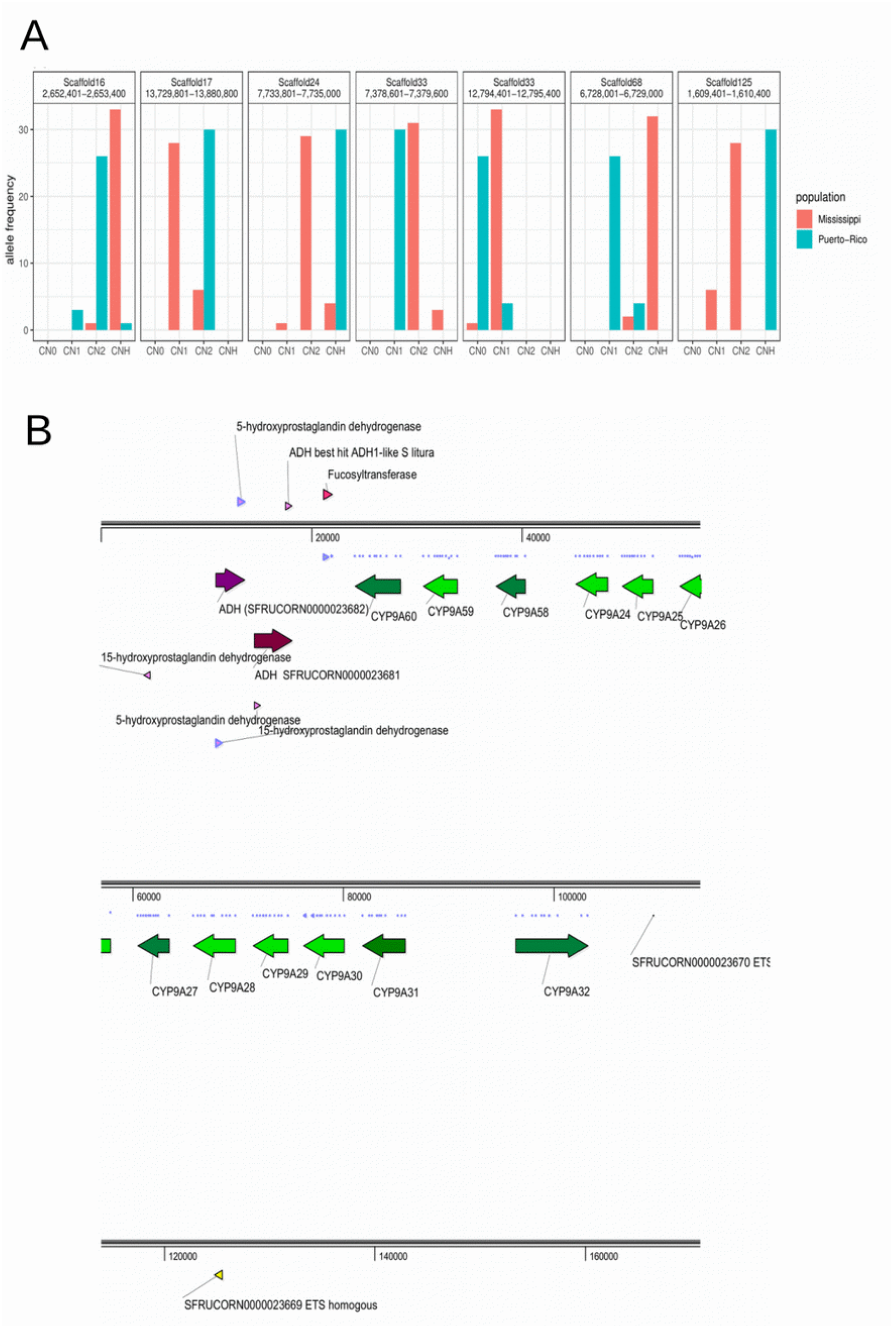
Positively selected CNV. A. Allele frequency of positively selected CNV. CN0, CN1, CN2, and CNH represent copy number equal to zero, one, two, and greater than two, respectively. B. The genes within the positively selected loci at the loci in scaffold 17. The purple and green colors represent ADH genes and P450 genes, respectively. The arrows indicate the direction of transcription.

The almost completely differentiated CNVs include a sequence at scaffold 24 as well (Fig 2A). In PR, the number of copy is always greater than three, while 88.2% of alleles in MS has one or two copies. Because multiple haplotypes exist both in PR and MS, it is unclear whether positive selection occurs specifically to PR because positive selection may also occur in MS in the way of reducing copy numbers. This CNV contains chitin deacetylase gene. Chitin deacetylase genes are widely used as a target of insecticides^35^, suggesting a possibility that CNV of this gene increased the resistance in PR. Chitin deacetylase is also associated with the response to *Bacillus thuringiensis* (Bt) in Lepidopteran *H. armigera* ^36^. PR has stronger resistance against Bt in corn than populations in the mainland of the USA^19,37,38^. Therefore, we propose a possibility that the increase in chitin deacetylase gene copy might be associated with the emergence of Bt-resistance in PR.

Scaffold 68 also contains almost completely differentiated CNV (Fig 2A). The copy number varies both in PR and MS. This CNV contains a gustatory receptor gene. The CNV of this gene is reported to be associated with the interaction with host-plants^10,33^. Thus, this CNV might be related to the adaptation to host-plants. However, the allelic differentiation between sfC and sfR is not observed at this loci. We did not identify any other protein-coding genes of known function from the remaining positively selected CNVs.

### Detoxification genes with CNV

Then, we tested if genes with CNVs are overrepresented in the lists of well-known detoxification genes, such CYP, esterase, Glutathione S-transferase (GST), UDP glucuronosyltransferases (UGT), and Oxidative stress genes. We observed that genes with CNV are significantly overrepresented in all the lists (FDR corrected p < 0.10) with the exception of CYP (FDR corrected p-value = 0.211) (Fig 3). This result strongly supports the association with detoxification and CNV.

**Figure 3.**
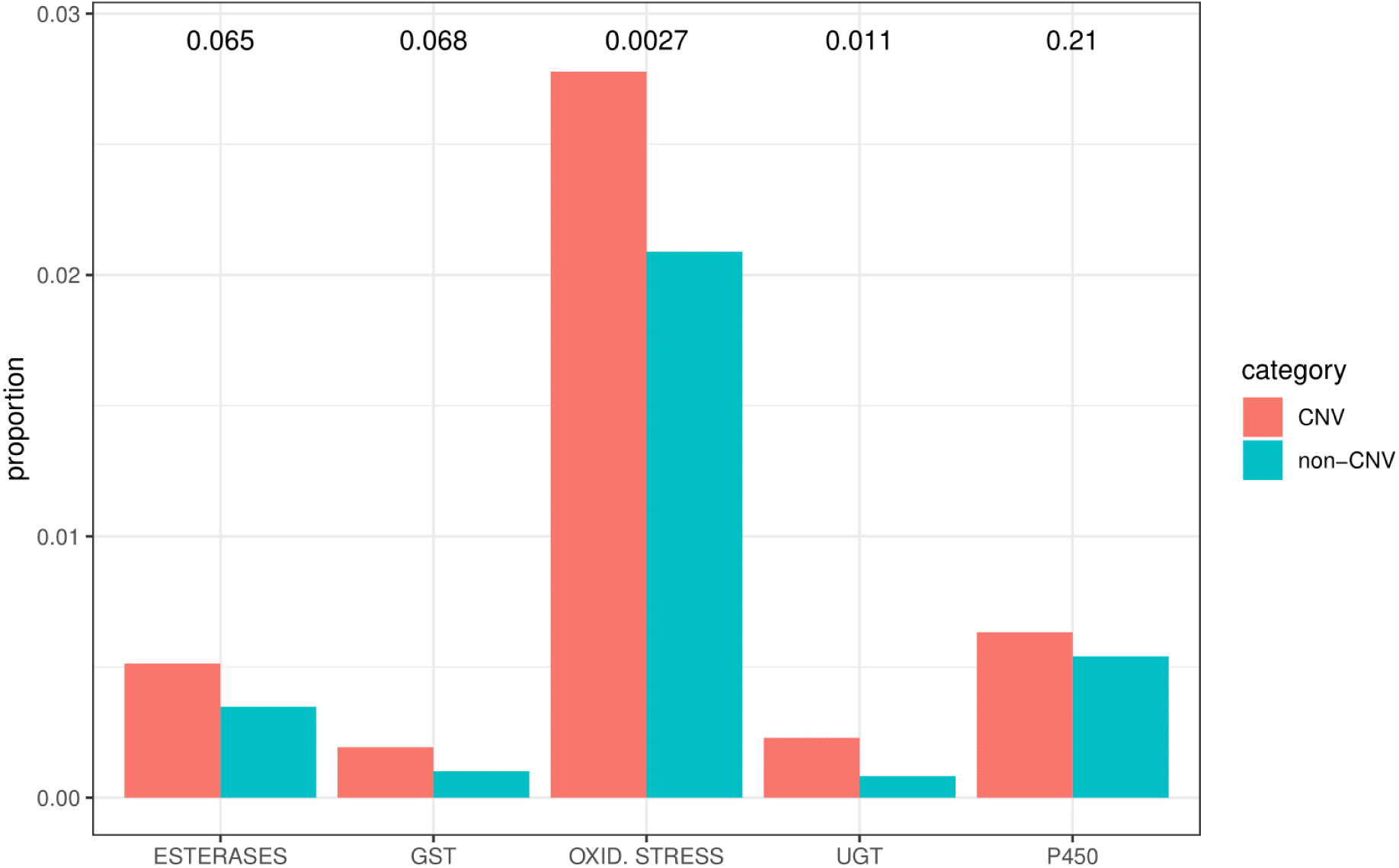
Testing the overrepresentation of detoxification genes with CNV. The bars indicate the proportion of detoxification genes in the genes with CNV or genes without CNV. The numbers above the bars indicate FDR-corrected p values.

We also performed gene ontology analysis to test the overrepresentation of other gene categories with CNVs using gene ontology analysis. In total, 31 gene ontology terms are overrepresented in the list of genes with CNVs (Table 2). These terms include anatomical structure development, developmental process, female gamete generation, and drug binding. This result shows that CNV might be functionally associated with other phenotypes than detoxification.

**Table 2.** The overrepresented gene ontology terms in the gene with CNV

| Category | description_Biological process | adjusted p value |
| --- | --- | --- |
| Biological Process | developmental process | 0.00184 |
|  | localization | 0.01494 |
|  | anatomical structure development | 0.04502 |
|  | establishment or maintenance of cell polarity | 0.06076 |
|  | multicellular organismal process | 0.06168 |
|  | multicellular organism development | 0.06377 |
|  | anatomical structure morphogenesis | 0.06938 |
|  | female gamete generation | 0.06938 |
| Cellular Component | chromosomal part | 0.05904 |
| Molecular Function | adenyl ribonucleotide binding | 0.00003 |
|  | adenyl nucleotide binding | 0.00003 |
|  | ATP binding | 0.00003 |
|  | drug binding | 0.00006 |
|  | purine ribonucleotide binding | 0.00010 |
|  | purine nucleotide binding | 0.00010 |
|  | purine ribonucleoside triphosphate binding | 0.00010 |
|  | ribonucleotide binding | 0.00012 |
|  | carbohydrate derivative binding | 0.00021 |
|  | anion binding | 0.00022 |
|  | ATPase activity | 0.00155 |
|  | small molecule binding | 0.00189 |
|  | nucleoside phosphate binding | 0.00264 |
|  | nucleotide binding | 0.00264 |
|  | binding | 0.00283 |
|  | ATPase activity, coupled | 0.00506 |
|  | ion binding | 0.01027 |
|  | protein binding | 0.01036 |
|  | hydrolase activity, acting on acid anhydrides | 0.06928 |
|  | hydrolase activity, acting on acid anhydrides, In phosphorus-containing anhydrides | 0.06928 |
|  | nucleoside-triphosphatase activity | 0.08469 |
|  | pyrophosphatase activity | 0.08535 |
|  | microtubule motor activity | 0.08535 |

### Beneficial effects of CNV

Then, we investigated the overall fitness effects of the observed CNVs. If the CNVs generate beneficial effects, then, the CNVs might be under the process of positive selection while yet to be fixed in a population. Alternatively, if the vast majority of CNVs have neutral or slightly deleterious effects, then the CNVs may evolve in an effectively neutral way. To test these possibilities, first, we compared the strength of selection between CNVs and SNVs with an assumption that exons are under stronger selective pressure than non-exonic sequences. The proportion of CNVs containing exonic sequences is 28.14% (11,720/41,645). This proportion is higher than that of SNVs (11.31%, 1,847,439/16,341,783, p < 2.2 × 10^−16^; Fisher’s exact test). The same trend was observed from mosquito, in which CNVs are overrepresented in gene-containing regions^8^.

This result can be interpreted by one of the following two possibilities. The first explanation is that the CNVs are experiencing stronger positive selection than SNVs because CNVs have a higher proportion of beneficial variants than SNV. The second explanation is that CNVs are under weaker purifying selection than SNVs because CNVs have weaker deleterious effects than SNV. According to population genetics theory, if purifying selection is weak, the proportion of rare alleles is increased because slightly deleterious variants still remain unpurged as rare variants in a population. To test the second possibility, we compared the proportion of singleton variants between CNVs and SNVs. In this case, we used the unfiltered CNV dataset. We observed that CNVs have a much lower proportion of singleton polymorphisms than SNVs (0.29% for CNV, 94.0% for SNV, < 2.2 × 10^−16^), suggesting that weaker purifying selection on CNVs is not supported. Instead, the first explanation, stronger positive selection on CNV, is a more likely scenario. The folded site frequency spectrum is very different between CNV and SNV(Fig 4A). Neutral or slightly deleterious effects are not likely to cause the observed pattern of folded site frequency spectrum of CNV, because, in this case, the proportion of singleton variants is expected to be highest among all site frequencies even in the presence of bottleneck^39^. Instead, the process of increasing derived allele frequency is a more likely explanation of this folded site frequency spectrum. The ancestry coefficient analysis shows that PR has a smaller number of CNV haplotypes than MS, supporting stronger positive selection in PR (Fig 4B).

**Figure 4.**
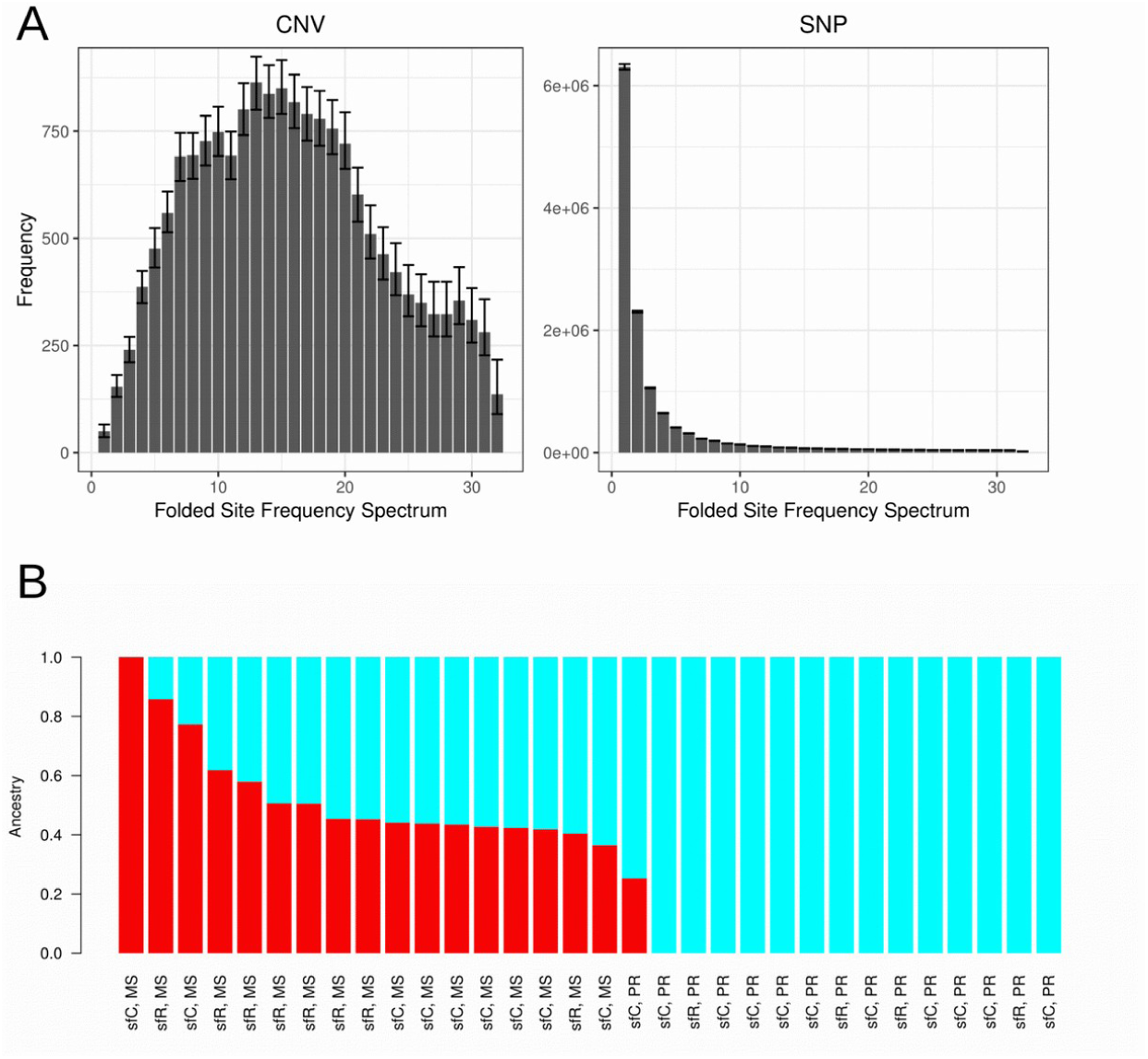
Selection on CNVs observed from PR. A. Folded site frequency spectrum at CNV and SNP. The error bars indicate 95% confidence intervals. B. The result of ancestry coefficient analysis.

## Discussion

In this study, we found that the adaptive evolution of detoxification genes by CNV increased the level of insecticide resistance in the fall armyworm. The analysis of the genomic CNVs between geographical populations with different levels of insecticide resistance and between strains with different host-plants enables us to distinguish the roles played by CNV between insecticide resistance and host-plant adaptation. We observed significant allelic differentiation of CNV between MS and PR, but not between host-plant strains. The loci with almost complete allelic differentiation between the geographic populations include a gene cluster containing CYP, which is one of the key players of the detoxification process, implying that this CNV potentially plays a crucial role in the increased level of insecticide resistance. The distribution of CNV across the genome is in line with beneficial effects. From these observations, we concluded that the CNV of detoxification genes increased insecticide resistance in PR.

This result shows that genes playing a key role in the increased insecticide resistance are different from that in the adaptation to host-plants. According to the “pre-adaptation hypothesis”, polyphagous insects have a greater ability to cope with toxins from a wide range of host-plants than monophagous ones. Thus, polyphagous insects have a higher potential to generate insecticide resistance^40,41^(but see ^42^). In this explanation, the emergence of insecticide resistance is no more than the re-use of existing detoxification genes that are adaptively evolved to host-plants. In polyphagous spider mites, genes that experienced host-plant adaptive evolution contribute to the increased level of insecticide resistance^43^, supporting this hypothesis. Our study proposes that even though detoxification machinery genes might largely overlap between host-plant and insecticide resistance, the key player for the insecticide resistance can evolve independently from the interaction with host-plants. In other words, the key player genes responsible for the insecticide resistance can be different from those for the adaptation to host-plants.

We showed that the genomic distribution of CNVs is generally in line with beneficial effects, implying prevalent positive selection on CNVs. Together with the result showing that detoxification genes are overrepresented in the genes with CNVs, these CNVs might increase the level of detoxification in a collective way. However, we wish to stress that not all of the beneficial CNVs will be fixed in a population. The probability of fixation of a beneficial mutation is 2*s*, where *s* is the selection coefficient of a beneficial allele. Thus, a very large proportion of mildly beneficial mutations will be lost eventually by genetic drift. In addition, the Hill-Robertson interference effect may hamper the fixation of adaptive CNV as well^44^. The increase of one adaptive CNV of detoxification genes may interfere with that of another adaptive CNV if these two CNVs are physically linked within a chromosome. Therefore, among the CNVs we observed, those with a strong beneficial effect will be fixed in a population. For this reason, we argue that the list of CNVs under positive selection should not be interpreted as positively *selected* CNV and that this gene list does not show the direction of adaptive evolution. Therefore, the observed CNV loci with almost complete differentiation (Fst > 0.8) are likely to have very strong beneficial effects.

The multiple inter-continental invasions of the fall armyworm is a serious global issue in agriculture. A marker-based study shows that these invasive populations were originated from the Florida-Caribbean region^45^. Therefore, the invasive population might carry the insecticide resistance CNVs, identified in this study. If this is true, invasive populations may have a higher degree of insecticide resistance than most of the native populations. In this case, the presence of insecticide resistance CNV might explain the success of the first invasion in West Africa. To test this possibility, assessing the existence of the insecticide resistance CNV in the invasive population and comparative studies on the insecticide resistance between native and invasive populations are urgently necessary.

This study is based on MS and PR, assuming that MS has a lower level of insecticide resistance than PR. The populations from Florida and the Caribbean have different mitochondrial haplotypes from the rest area of North and South American continents^46,47^, possibly due to the physical barrier effect of the Appalachian mountain range^48^ and the limited genetic exchanges through the Lesser Antilles^47^.This grouping implies that the long-term seasonal migration of the fall armyworm is not sufficient for genetic exchanges between Puerto Rico and the Western area of the Appalachian mountain range to the observable extent. Therefore, we assumed that genes responsible for the insecticide resistance of fall armyworms in Puerto Rico were not introgressed in Mississippi’s populations. However, we acknowledge the possibility that this MS might also have an increased level of insecticide resistance for an unknown reason. If this possibility is true, and if the gene flow between PR and MS is responsible for the increased level of insecticide resistance in MS, then CNVs responsible for the increased insecticide resistance might remain unidentified in this study. To elucidate this issue, a comparative study on the level of insecticide resistance in a wide range of geographical populations is necessary.

## Conclusions

We showed that positive selection on the CNV of detoxification genes is related to the insecticide resistance of PR of the fall armyworm using genome-scan. This *in silico* approach can be complemented by functional genetics studies to verify the role of each positively selected loci CNV in the increased level of insecticide resistance. For example, CRISPR/Cas9^49^ or RNAi^50^ for each CNVs can be used for this purpose. Importantly, the association between CNV and gene expression level can be useful to verify the role of CNV in the differential levels of insecticide resistance. All these approaches enable us to obtain a more comprehensive and accurate knowledge of the insecticide resistance, which will ultimately lead to establishing realistic and effective ways of insect pest control.

## Methods

### data

Pooled high molecular weight genomic DNA was extracted from 10 pupae of sfC, which was seeded from Guadeloupe and have been raised at our insectarium. Then, Dovetail™ Hi-C library was constructed from this strain, followed by Illumina Hiseq-X sequencing. Then, we used Hi-Rise™ software^24^ to perform scaffolding from a PacBio genome assembly^22^, which was generated from individuals from the same insectarium. Then, the gene annotation was also transferred from this PacBio genome assembly to newly scaffoled-assembly. The correctness of the genome assembly was assessed using BUSCO^26^.

The resequencing data from the MS was obtained from NCBI SRA (PRJNA494340), which was generated for the fall armyworm genome project^10^ and speciation study^22^. For PR, we collected larvae samples at Santa Isabella (see^38^ for more detail). The larvae were raised to adults, and gDNA was extracted from thorax using Promega Wizard® Genomic DNA Purification Kit. Then, 150bp pair-end Hiseq-4000 sequencing was performed from 15 individuals. These resequencing data have approximately 20X coverage per individual. The adapter sequences were removed from the adapterremoval^51^. To identify strains, we mapped the Illumina reads against mitochondrial genomes (NCBI: KM362176) using bowtie2^52^, followed by performing variant calling together with individuals from MS using samtools^53^. The strain of individuals from MS was already identified from mitochondrial CO1 in our previous study^10^. We performed principal component analysis to observe the grouping according to the strain using picard^54^. For nuclear mapping, first, the reads were mapped against the reference genome using bowtie2^52^ with –very-sensitive-local preset option. Potential optical or PCR duplicates were removed using picard tools^54^.

### variant identification

Variant calling was performed using GATK-4.0.11.0 package^27^.First, we performed haplotype calling, and resulting gvcf files were merged using this package. Then, variants were called, and only SNPs were extracted among all variants. The number of identified SNPs is 66,529,611. The SNPs were filtered out if QualByDepth is less than 2, or FisherStrand score is higher than 60, or RMSMappingQuality is less than 40, or MappingQualityRankSumTest score is less than −12.5, or ReadPosRankSumTest is less than −8. The number of remaining SNP is 16,341,783.

CNVs were inferred from the bam files, which were generated by the mapping of Illumina reads against reference genome assembly, using CNVcaller^28^, with 100bp window size. We filtered out CNVs if the minor allele frequency is less than 0.1 to reduce false positives, based on the assumption that the probability of having false positives at the same genomic locus from multiple samples is very low. In other words, we analyzed the CNVs only if CNV was observed from at least four individuals out of total 32 (corresponding to allele frequency equal to 4/32 = 0.125) within 100bp intervals.

### Population genetics analysis

The principal component analysis was performed using picard^54^ after converting the vcf file to plink format using vcftools^55^ and picard^54^. Pairwise Weir and Cockerham’s Fst^56^ was calculated using VCFtools^55^. The gene ontology analysis was performed using BinGO^57^. The ancestry coefficient analysis was performed using admixture^58^.

### Data availability

The resequencing data in this study is available in NCBI SRA (PRJNA494340, PRJNA577869), and the reference genome assembly is available in BIPAA (www.bipaa.genouest.org/sp/spodoptera_frugiperda).

## Author Contributions

KN: Conceptualization; Formal Analysis; Funding Acquisition, Investigation, Project Administration, Writing the paper; SG: Preparation samples for sequencing; FH: Manual gene annotation; CAB, SH: Acquisition of samples for sequencing; AB, FL: Generation of the reference genome, automatic gene annotation; NN, EA: The acquisition of Illumina sequencing data from the population in Mississippi. All co-authors participated in the writing of this paper, and they approved it.

## Acknowledgments

We acknowledge Yutao Xiao (CAAS) for the discussion about this paper. We also acknowledge Claire Lemaitre (INRIA), who helped us to identify CNV.

## References

1. Palumbi, S. R. Humans as the World’s Greatest Evolutionary Force. Science 293, 1786–1790 (2001).

2. Alphey, N. & Bonsall, M. B. Genetics-based methods for agricultural insect pest management. Agricultural and forest entomology 20, 131–140 (2018).

3. David, J.-P., Ismail, H. M., Chandor-Proust, A. & Paine, M. J. I. Role of cytochrome P450s in insecticide resistance: impact on the control of mosquito-borne diseases and use of insecticides on Earth. Philos Trans R Soc Lond B Biol Sci 368, (2013).

4. Montella, I. R., Schama, R. & Valle, D. The classification of esterases: an important gene family involved in insecticide resistance--a review. Mem. Inst. Oswaldo Cruz 107, 437–449 (2012).

5. Li, X., Shi, H., Gao, X. & Liang, P. Characterization of UDP-glucuronosyltransferase genes and their possible roles in multi-insecticide resistance in Plutella xylostella (L.). Pest Manag. Sci. 74, 695–704 (2018).

6. Oliver, S. V. & Brooke, B. D. The Role of Oxidative Stress in the Longevity and Insecticide Resistance Phenotype of the Major Malaria Vectors Anopheles arabiensis and Anopheles funestus. PLoS One 11, (2016).

7. Ranson, H. et al. Evolution of Supergene Families Associated with Insecticide Resistance. Science 298, 179–181 (2002).

8. Lucas, E. R. et al. Whole-genome sequencing reveals high complexity of copy number variation at insecticide resistance loci in malaria mosquitoes. Genome Res. 29, 1250–1261 (2019).

9. Gong, J. et al. Genome-wide patterns of copy number variations in Spodoptera litura. Genomics (2018) doi:10.1016/j.ygeno.2018.08.002.

10. Gouin, A. et al. Two genomes of highly polyphagous lepidopteran pests (Spodoptera frugiperda, Noctuidae) with different host-plant ranges. Scientific Reports 7, 11816 (2017).

11. Weetman, D., Djogbenou, L. S. & Lucas, E. Copy number variation (CNV) and insecticide resistance in mosquitoes: evolving knowledge or an evolving problem? Curr Opin Insect Sci 27, 82–88 (2018).

12. War, A. R. et al. Plant defence against herbivory and insect adaptations. AoB PLANTS 10, (2018).

13. Goergen, G., Kumar, P. L., Sankung, S. B., Togola, A. & Tamò, M. First Report of Outbreaks of the Fall Armyworm Spodoptera frugiperda (J E Smith) (Lepidoptera, Noctuidae), a New Alien Invasive Pest in West and Central Africa. PLOS ONE 11, e0165632 (2016).

14. Kalleshwaraswamy, C. M. et al. First report of the fall armyworm, Spodoptera frugiperda (JE Smith)(Lepidoptera: Noctuidae), an alien invasive pest on maize in India. Pest Management in Horticultural Ecosystems 24, 23–29 (2018).

15. Pashley, D. P. Host-associated Genetic Differentiation in Fall Armyworm (Lepidoptera: Noctuidae): a Sibling Species Complex? Ann Entomol Soc Am 79, 898–904 (1986).

16. Pashley, D. P. Host-associated differentiation in armyworms (Lepidoptera: Noctuidae): An allozymic and mtDNA perspective. Electrophoretic studies on agricultural pests (1989).

17. Dumas, P. et al. Spodoptera frugiperda (Lepidoptera: Noctuidae) host-plant variants: two host strains or two distinct species? Genetica 143, 305–316 (2015).

18. Orsucci, M. et al. Transcriptional plasticity evolution in two strains of Spodoptera frugiperda (Lepidoptera: Noctuidae) feeding on alternative host-plants. bioRxiv 263186 (2018) doi:10.1101/263186.

19. Zhu, Y. C. et al. Evidence of multiple/cross resistance to Bt and organophosphate insecticides in Puerto Rico population of the fall armyworm, Spodoptera frugiperda. Pestic Biochem Physiol 122, 15–21 (2015).

20. Gutiérrez-Moreno, R. et al. Field-Evolved Resistance of the Fall Armyworm (Lepidoptera: Noctuidae) to Synthetic Insecticides in Puerto Rico and Mexico. J Econ Entomol 112, 792–802 (2019).

21. Belay, D. K., Huckaba, R. M. & Foster, J. E. Susceptibility of the fall armyworm, Spodoptera frugiperda (Lepidoptera: Noctuidae), at Santa Isabel, Puerto Rico, to different insecticides. Florida Entomologist 95, 476–479 (2012).

22. Nam, K. et al. Divergent selection causes whole genome differentiation without physical linkage among the targets in Spodoptera frugiperda (Noctuidae). bioRxiv 452870 (2018) doi:10.1101/452870.

23. Lieberman-Aiden, E. et al. Comprehensive Mapping of Long-Range Interactions Reveals Folding Principles of the Human Genome. Science 326, 289–293 (2009).

24. Putnam, N. H. et al. Chromosome-scale shotgun assembly using an in vitro method for long-range linkage. Genome Res 26, 342–350 (2016).

25. Liu, H. et al. Chromosome level draft genomes of the fall armyworm, Spodoptera frugiperda (Lepidoptera: Noctuidae), an alien invasive pest in China. bioRxiv 671560 (2019) doi:10.1101/671560.

26. Simão, F. A., Waterhouse, R. M., Ioannidis, P., Kriventseva, E. V. & Zdobnov, E. M. BUSCO: assessing genome assembly and annotation completeness with single-copy orthologs. Bioinformatics 31, 3210–3212 (2015).

27. McKenna, A. et al. The Genome Analysis Toolkit: A MapReduce framework for analyzing next-generation DNA sequencing data. Genome Res 20, 1297–1303 (2010).

28. Wang, X. et al. CNVcaller: highly efficient and widely applicable software for detecting copy number variations in large populations. Gigascience 6, (2017).

29. Nagoshi, R. N. The fall armyworm Triosephosphate Isomerase (Tpi) gene as a marker of strain identity and interstrain mating. Ann Entomol Soc Am 103, 283–292 (2010).

30. d’Alençon, E. et al. Extensive synteny conservation of holocentric chromosomes in Lepidoptera despite high rates of local genome rearrangements. PNAS 107, 7680–7685 (2010).

31. Giraudo, M. et al. Cytochrome P450s from the fall armyworm (Spodoptera frugiperda): responses to plant allelochemicals and pesticides. Insect Mol. Biol. 24, 115–128 (2015).

32. Hu, B. et al. The expression of Spodoptera exigua P450 and UGT genes: tissue specificity and response to insecticides. Insect Sci. 26, 199–216 (2019).

33. Cheng, T. et al. Genomic adaptation to polyphagy and insecticides in a major East Asian noctuid pest. Nature Ecology & Evolution 1, 1747–1756 (2017).

34. Uchibori-Asano, M. et al. Genome-wide Identification of Tebufenozide Resistant Genes in the smaller tea tortrix, Adoxophyes honmai (Lepidoptera: Tortricidae). Sci Rep 9, 1–12 (2019).

35. Zhao, Y., Park, R.-D. & Muzzarelli, R. A. A. Chitin Deacetylases: Properties and Applications. Marine Drugs 8, 24–46 (2010).

36. Han, G., Li, X., Zhang, T., Zhu, X. & Li, J. Cloning and Tissue-Specific Expression of a Chitin Deacetylase Gene from Helicoverpa armigera (Lepidoptera: Noctuidae) and Its Response to Bacillus thuringiensis. J. Insect Sci. 15, (2015).

37. Storer, N. P., Kubiszak, M. E., Ed King, J., Thompson, G. D. & Santos, A. C. Status of resistance to Bt maize in Spodoptera frugiperda: lessons from Puerto Rico. J. Invertebr. Pathol. 110, 294–300 (2012).

38. Blanco, C. A. et al. Susceptibility of isofamilies of Spodoptera frugiperda (Lepidoptera: Noctuidae) to Cry1Ac and Cry1Fa proteins of Bacillus thuringiensis. Southwestern Entomologist 35, 409–416 (2010).

39. Watterson, G. A. Allele frequencies after a bottleneck. Theoretical Population Biology 26, 387–407 (1984).

40. Gordon, H. T. Nutritional Factors in Insect Resistance to Chemicals. Annual Review of Entomology 6, 27–54 (1961).

41. Alyokhin, A. & Chen, Y. H. Adaptation to toxic hosts as a factor in the evolution of insecticide resistance. Current Opinion in Insect Science 21, 33–38 (2017).

42. Dermauw, W., Pym, A., Bass, C., Van Leeuwen, T. & Feyereisen, R. Does host plant adaptation lead to pesticide resistance in generalist herbivores? Current Opinion in Insect Science 26, 25–33 (2018).

43. Dermauw, W. et al. A link between host plant adaptation and pesticide resistance in the polyphagous spider mite Tetranychus urticae. PNAS 110, E113–E122 (2013).

44. The Hill–Robertson effect: evolutionary consequences of weak selection and linkage in finite populations | Heredity. https://www.nature.com/articles/6801059.

45. Nagoshi, R. N. et al. Analysis of strain distribution, migratory potential, and invasion history of fall armyworm populations in northern Sub-Saharan Africa. Scientific Reports 8, 3710 (2018).

46. Nagoshi, R. N. et al. Haplotype Profile Comparisons Between Spodoptera frugiperda (Lepidoptera: Noctuidae) Populations From Mexico With Those From Puerto Rico, South America, and the United States and Their Implications to Migratory Behavior. J Econ Entomol 108, 135–144 (2015).

47. Nagoshi, R. N. et al. Fall armyworm migration across the Lesser Antilles and the potential for genetic exchanges between North and South American populations. PLOS ONE 12, e0171743 (2017).

48. Nagoshi, R. N., Meagher, R. L. & Hay-Roe, M. Inferring the annual migration patterns of fall armyworm (Lepidoptera: Noctuidae) in the United States from mitochondrial haplotypes. Ecol Evol 2, 1458–1467 (2012).

49. Zhang, L. & Reed, R. D. A Practical Guide to CRISPR/Cas9 Genome Editing in Lepidoptera. in Diversity and Evolution of Butterfly Wing Patterns: An Integrative Approach (eds. Sekimura, T. & Nijhout, H. F.) 155–172 (Springer Singapore, 2017). doi:10.1007/978-981-10-4956-9_8.

50. Terenius, O. et al. RNA interference in Lepidoptera: an overview of successful and unsuccessful studies and implications for experimental design. J. Insect Physiol. 57, 231–245 (2011).

51. Schubert, M., Lindgreen, S. & Orlando, L. AdapterRemoval v2: rapid adapter trimming, identification, and read merging. BMC Research Notes 9, 88 (2016).

52. Langmead, B. & Salzberg, S. L. Fast gapped-read alignment with Bowtie 2. Nat. Methods 9, 357–359 (2012).

53. Li, H. et al. The sequence alignment/map format and SAMtools. Bioinformatics 25, 2078–2079 (2009).

54. picard: A set of command line tools (in Java) for manipulating high-throughput sequencing (HTS) data and formats such as SAM/BAM/CRAM and VCF. (Broad Institute, 2018).

55. Danecek, P. et al. The variant call format and VCFtools. Bioinformatics 27, 2156–2158 (2011).

56. Weir, B. S. & Cockerham, C. C. Estimating F-statistics for the analysis of population structure. Evolution 38, 1358–1370 (1984).

57. Maere, S., Heymans, K. & Kuiper, M. BiNGO: a Cytoscape plugin to assess overrepresentation of Gene Ontology categories in Biological Networks. Bioinformatics 21, 3448–3449 (2005).

58. Alexander, D. H., Novembre, J. & Lange, K. Fast model-based estimation of ancestry in unrelated individuals. Genome Res 19, 1655–1664 (2009).

