## Supplemental for "Adaptation by copy number variation increases insecticide resistance in fall armyworms"

Supplementary information

Contents:

Supplementary Figure 1. The distribution of scaffold lengths.
Supplementary Figure 2. The result of PCA to identify strains
Supplementary Figure 3. The coverage of mapped reads for each sample.
Supplementary Figure 4. The level of genetic differentiation of autosomes and Z chromosomes in the absence and presence of hybridization.

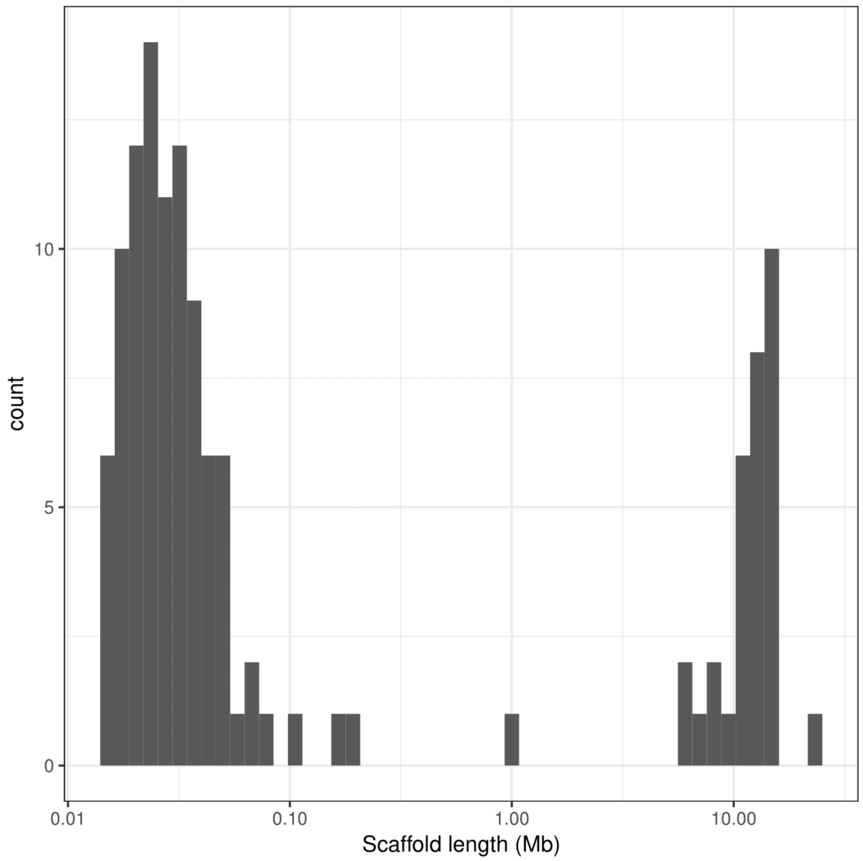


Supplementary Figure 1. **The distribution of scaffold lengths.** It shows a bimodal distribution, and the number of scaffolds at the left mode is 31, which is the same as the chromosome number in the fall armyworm.

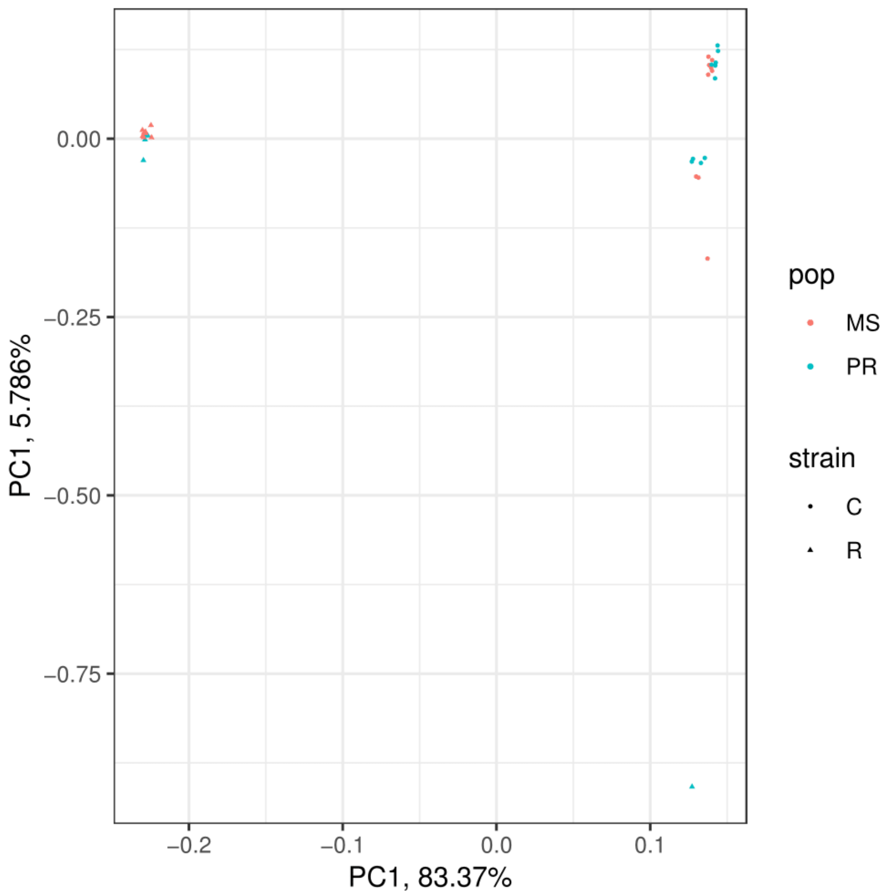


Supplementary Figure 2. The result of PCA to identify strains. C and R indicate sfC and sfR, respectively. MS and PR indicate a population from Mississippi and Puerto Rico, respectively.

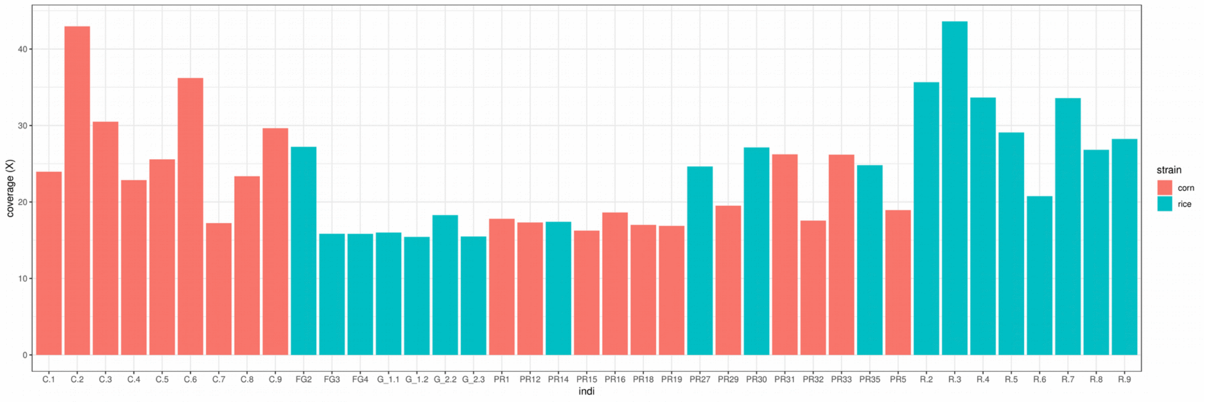


Supplementary Figure 3. The coverage of mapped reads for each sample.

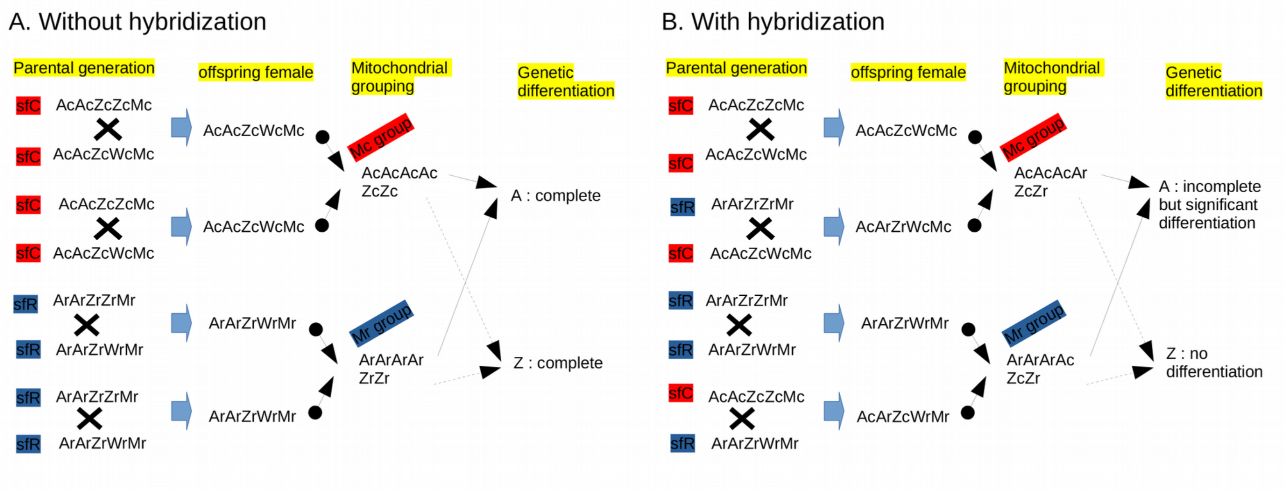


Supplementary Figure 4. **The level of genetic differentiation of autosomes and Z chromosomes in the absence and presence of hybridization**. A, Z, W, and M indicate Autosome, Z chromosome, W chromosome, and Mitochondrion, respectively. The small capital of c and r indicate the genotypes of sfC and sfR, respectively. For example, Ac indicates an Autosomal sfC genotype, and Ar indicates an Autosomal sfR genotype. We assume that each parental pair generate one female because we analyzed only females in this study. A. Without hybridization. In the parental generation, two pairs of sfC parents mate ( AcAcZcWcMc females and AcAcZcZcMc males). In the following generation, two daughters have the same genotype, AcAcZcWcMc. In the parental generation, two pairs of sfR parents mate (ArArZrWrMr females and ArArZrZrMr males) and two generated daughters have ArArZrWrMr genotype. When we group these genotypes according to the mitochondrial sequences, in the Mc group, there are four Ac genotypes and two Zc genotypes. In the Mr group, there are four Ar genotypes and two Zr genotypes. When we calculate the level of genetic differentiation (e.g., Fst) between the Mc and Mr groups, both A and Z will show complete genetic differentiation because the genotype of A in Mc is always Ac and that in Mr is always Ar and because the genotype of Z in Mc is always Zc and that in Mr is always Zr. B. With hybridization. In the parental generation, one pair of sfC parents mate, and in the following generation, a generated daughter will have the AcAcZcWcMc genotype. In another parental pair, sfR father and sfC mother mate, the generated daughter will have the AcArZrWcMc genotype. In another parental pair, both parents are sfR. Thus the generated genotype is ArArZrWrMr. In the last parental pair, sfC father and sfR mother mate, then the generated daughter will have the AcArZcWrMr genotype. When we classify these genotypes according to the mitochondrial genotype, in the Mc group, the numbers of Ac, Ar, Zc, and Zr are three, one, one, and one, respectively. In the Mr group, the numbers of Ac, Ar, Zc, Zr are one, three, one, and one, respectively. When we calculate the level of genetic differentiation between Mc and Mr groups, A will show significant genetic differentiation because of unequal number of Ac and Ar, while the level of genetic differentiation will show incomplete differentiation because one out of four As will show a different genotype from the other three (There are one Ac and three Ar in Mc group, and there are three Ar and one Ac in Mr group). Z will not show a significant genetic differentiation because the number of Zc and Zr is the same in both Mc and Mr group. As a consequence, in the presence of hybridization, A will show a higher level of genetic differentiation than Z in females.
